# Fecal microbiota transplantation reconstructs the gut microbiota of septic mice and protects the intestinal mucosal barrier

**DOI:** 10.1101/2020.06.22.164541

**Authors:** Xiaowei Gai, Huawei Wang, Yaqing Li, Haotian Zhao, Cong He, Zihui Wang, Heling Zhao

**Affiliations:** Graduate school of Hebei Medical University; Hebei General Hospital; Qinhuangdao cerebrovascular disease hospital

**Keywords:** sepsis, fecal microbiota transplantation, intestinal mucosal barrier, gut microbiota, critical care

## Abstract

The gastrointestinal (GI) tract has long been hypothesized to play an integral role in the pathophysiology of sepsis, and gut microbiota (GM) dysbiosis may be the key factor. Previous studies has confirmed that microbiome is markedly altered in critical illness. We aimed to confirm the existence of gut microbiota imbalance in the early stage of sepsis, observe the effect of fecal microbiota transplantation (FMT) on sepsis, and explore whether FMT can reconstruct the GM of septic mice and restore its protective function on the intestinal mucosal barrier. Through the study of flora, mucus layer, tight junction, immune barrier, and short-chain fatty acid changes in septic mice and fecal microbiota transplanted mice, we found that GM imbalance exists early in sepsis. FMT can improve morbidity and effectively reduce mortality in septic mice. After the fecal bacteria were transplanted, the abundance and diversity of the gut flora were restored, and the microbial characteristics of the donors changed. FMT can effectively reduce epithelial cell apoptosis, improve the composition of the mucus layer, upregulate the expression of tight junction proteins, and reduce intestinal permeability and the inflammatory response, thus protecting the intestinal barrier function. After FMT, Lachnospiraceae contributes the most to intestinal protection through enhancement of the L-lysine fermentation pathway, resulting in the production of acetate and butanoate, and may be the key bacteria for short-chain fatty acid metabolism and FMT success.

## Introduction

Sepsis continues to be the leading cause of mortality in the intensive care unit^1, 2^. There were more than 1 million sepsis-related deaths in 2015 in China^3^. In 2017, an estimated 48.9 million incident cases of sepsis were recorded worldwide, with 11.0 million sepsis-related deaths reported, comprising 19.7% of global deaths^4^. The World Health Organization has recognized sepsis as a global health priority^5^. Despite significant advancement in our understanding of the pathophysiology of sepsis, treatment of sepsis is limited to antibiotics, aggressive fluid resuscitation, vasopressor administration, and supportive care, and no targeted therapeutics for sepsis are approved for usage in patients^6^.

Sepsis is a life-threatening organ dysfunction caused by a dysregulated host response to infection^5–7^. The syndrome can be induced by a wide variety of microbes and, by definition, involves a maladaptive response to a pathogen^5^. The gastrointestinal (GI) tract has long been hypothesized to play an integral role in the pathophysiology of sepsis, by both driving and perpetuating multiple organ dysfunction^8, 9^. The original concept of gut-derived sepsis proposed that the altered inflammatory milieu induced by overwhelming infection leads to intestinal hyperpermeability, allowing luminal contents, including intact microbes, and microbial products, to escape their natural environment where they can cause either local or distant injury^1, 5, 6, 10^.

The gut microbiome plays a crucial role not only in GI health but also in overall immune development and host health. The gut microbiome contains 40 trillion microorganisms, a similar number of cells as found in the human host. This includes up to 1000 different microbial species and 100 times more bacterial genes than human genes^6, 10–13^. The symbiotic relationship between microbiota and the host is mutually beneficial^14^. The host provides an important habitat and nutrients for the microbiome, and the gut microbiota (GM) support the development of the metabolic system and the maturation of the intestinal immune system by providing beneficial nutrients, for example, by the synthesis of vitamins and short-chain fatty acids (SCFAs)^15^. The destruction of the balance between the gut microbiome and the host is called dysbiosis. The microbiome is markedly altered in critical illness. Microbial diversity is diminished within 6 h of admission to the intensive care unit, and this lack of diversity has been associated with poor outcomes in critically ill patients^5^.

Considering GM dysbiosis is one of the most important factors that can lead to pathological bacterial translocation and systemic infection, it may be feasible to develop novel therapeutic strategies against gut-derived sepsis by modulating the microbiota^16^. More than 90% of the commensal organisms may be lost during the early stage of critical illness, making it nearly impossible that a single or several probiotic species would be able to completely replenish the diversity of the GM without intervention. Transfer of healthy donor feces containing thousands of microbial species, termed FMT, facilitates replenishment of diminished commensal bacteria and may guide the patient’s microbiota toward a healthy state^16^. Fecal microbiota transplantation has been demonstrated to be remarkably successful in the treatment of recurrent *Clostridium difficile* infection, with a 92% response rate to treatment, and is also increasingly being used for dysbiosis caused by other intestinal pathologies, such as inflammatory bowel disease^12, 17^. Yet, FMT is scarcely used in the treatment of septic patients, as antibiotic therapy is frequently used in these cases and continuation of FMT therapy could adversely influence the remodeling of intestinal microbiota^16^. Recently, the use of FMT has been reported in septic patients with MODS and non-*C.difficile* diarrhea, presenting as refractory to standard medical management. At 2-3 weeks post-FMT, the patients showed resolution of their diarrhea and significant decreases in blood levels of inflammatory mediators, such as TNF-α, interleukin (IL)-1β, IL-6, and C-reactive protein^11, 16^. Following FMT, stool microbiota in these patients showed marked alterations resembling the microbiota composition of the donors, with an increase in Firmicutes and a reduction in Proteobacteria. Even though the evidence is limited to a series of case reports, the improved clinical outcomes in these patients following FMT are promising^16, 18^. However, during sepsis, the exact mechanism of action for the use of FMT on the intestine is still unknown^19^.

## Materials and methods

### Mice

The study included 6-8-week-old male clean-grade C57BL/6 mice weighing between 20-25g (Beijing Vital River Laboratory Animal Technology Co.). The variation in intestinal microbiota composition between inbred C57BL/6 mice derived from one commercial vendor is known to be highly limited^20^. Breeding conditions included a 12/12-hour light/dark cycle at a room temperature of 20-25°C. All mice were given free access to food and water. This experiment was approved by the Committee on the Ethics of Animal Experiments of the Hebei General Hospital.

### Sepsis model

Sepsis was induced by cecal ligation and puncture (CLP) as a previously described^21, 22^. Mice were anesthetized using an intraperitoneal injection at 50 mg/kg of 2% sodium pentobarbital. Briefly, a 1.5 cm midline laparotomy was performed under aseptic conditions to expose the cecum. A single through-and-through puncture was performed by a 22-gauge needle between the ligation site and the end of the cecum, and then a small amount of fecal material was extruded through the puncture. The cecum was repositioned into the peritoneal cavity, and the peritoneum and skin were closed in layers. Sham mice were treated identically except the cecum was neither ligated nor punctured. All mice received 1.0 ml normal saline subcutaneously after the surgery to compensate for fluid loss. Mice were euthanized at 12, 24, and 48 h following CLP for acute studies (Sham: n = 6 per group; CLP: n = 10/11 per group; FMT: n = 10/11 per group).

### Fecal microbiota transplantation

Fresh feces was collected from 10 healthy C57BL/6 mice, that were from the same strain of the recipient, homogenized in 10 ml of sterile phosphate-buffered saline (PBS) and centrifuged for 30 sec at 3,000 rpm, 4 °C, to pellet the particulate matter. The optical density (OD) value of the supernatant slurry was checked to calculate the concentration of total bacteria (OD = 0.5 represents 10^8^ cells). For each mouse, 1×10^8^ bacterial cells (1×10^9.8^ bacterial cells represents the sum of the total bacterial population within 2 g of cecal contents) were centrifuged for 5 min at 12,000 rpm, at 4 °C, and them bacterial pellets were resuspended in 0.2 ml PBS and gavaged into each mouse one day at a time^23, 24^. For acute studies, mice in the FMT group received a single dose of fecal microbiota just prior to cecal ligation and puncture for the first day^22^. The CLP and Sham group mice were gavaged with 0.2 ml PBS one day at a time. All mice were treated at the same time. For the survival studies, mice in the FMT group were treated with fecal microbiota once a day for the first 3 days, and the other mice were treated with PBS daily for 3 days.

### Serum IL-6, IL-10, and TNF-α analysis

Enzyme-linked immunosorbent assay was used to determine the concentrations of TNF-α (Lot: RXQJYSXLD4, Elabscience, China), IL-6 (Lot: 449268JEI6, Elabscience, China), and IL-10 (Lot: C8QEY2KAVE, Elabscience, China) in serum (Sham group n = 6 per group; CLP n = 6 per group; FMT n = 6 per group) according to the manufacturer’s instructions. Serum was collected after centrifuging blood for 10 min at 4 °C and 3,500 rpm and stored at −80°C until the assay was performed.

### Sample processing for animal experiments

After the mice were sacrificed, their GI tracts were quickly removed. The colons were gently separated, by cutting at the cecum-colon junction and rectum, and immediately preserved in Carnoy’s fixative (dry methanol: chloroform: glacial acetic acid in the ratio 60:30:10)^25, 26^. The Carnoy’s fixative was made fresh with anhydrous methanol, chloroform and glacial acetic acid. The colons were fixed in Carnoy’s solution for 3 h followed by transfer to fresh Carnoy’s solution for 2-3 h. The colons were then washed in dry methanol for 2 h, placed in cassettes and stored in fresh dry methanol at 4°C until further use. Cecal contents from each animal were divided into replicates, and each distal end was instantly flash-frozen in liquid nitrogen and then stored at −80°C until further use.

### Morphological examination and histological analysis

The colonic sections were stained with hematoxylin/eosin (HE) and examined under a phase-contrast microscope for morphological characteristics. The histological damage was scored using previously published criteria^23^, including extent of destruction of normal epithelial architecture, presence and degree of inflammatory infiltration, presence of edema, extent of vascular dilatation and congestion, and presence or absence of goblet cell depletion, presence or absence of crypt abscesses. To observe the thickness of the mucus layer, we stained with Alcian, and observed under a DP80 microscope (Olympus, Japan).

### Immunohistochemistry and immunofluorescence

Samples were fixed in Carnoy’s fixative, embedded in paraffin, and cut into sections (5μm thickness). Sections were mounted onto polylysine-coated slides, deparaffinized, rehydrated and placed in a 3% citrate buffer to repair antigens. After being pretreated with 3% H2O2 for 30 min, the sections were blocked with goat serum for 20 min. Sections were incubated overnight at 4°C with a rabbit polyclonal anti-mouse blocking antibody (MUC2, ab90007, Abcam Ltd; Occludin, ab168986, Abcam Ltd; caspase 3, ab44976, Abcam Ltd). The sections were washed with PBS and incubated with a secondary antibody for 30 min, rewashed, and incubated with peroxidase-conjugated streptavidin for 30 min. DAB developed, hematoxylin counterstained, dehydrated, and mounted. The secondary antibody of MUC 2 was prepared in 0.5% Triton X-100 PBS buffer, and the Alexa Fluor™ 488 donkey anti-rabbit antibody was diluted 1:1000. The specimen was incubated in this solution at 37°C for 1 h. We aspirated the secondary antibody and rinsed the specimen three times in PBS for 5 min each and covered the specimen with a DAPI coverslip.

### Transmission electron miceroscopy (TEM)

For TEM, two 0.5×0.5 cm mini-segments of intestinal tissue from each group were excised and placed in fixative for TEM at 4°C for 2-4 h. The segments were washed in 0.1 M PBS three times for 15 min each time, post fixed in 1% osmium tetroxide in PBS, dehydrated in a graduated series of ethanol solution, and embeded by baking at 60°C oven for 48 h. Samples were cut into 60-80 nm sections, and stained with uranyl acetate and lead citrate. The sections were analyzed by electronic microscopy (HT7700 TEM; Hitachi Inc., Tokyo, Japan).

### Western blot

The snap-frozen tissues were subjected to homogenization in 250μl of lysis buffer as previously described. Samples (of 30 μg of protein for each condition) were transferred onto PVDF membranes and then incubated with antibodies^24^. The following were used as primary antibodies: caspase 3 (ab323519, Abcam), myeloid differentiation factor 88 (MyD88) (SC74532, Santa Cruze), toll-like receptor 4 (TLR4) (AF7017, Affinity), ZO-1 (AF 5145, Affinity), occludin (DF7504, Affinity), and NF-κB(ab16502, Abcam). Immunoreactive bands were revealed using a 1:10,000 dilution of secondary antibody conjugated to horseradish peroxidase (goat anti-rabbit IgG, BE0101, Bioeasy; goat anti-mice IgG, BE0102, Bioeasy). The blots were re-probed with antibodies against β-actin (EASYBIO) and GAPDH (Bioworld) to ensure equal loading and transfer of proteins. All critical blots and immunoprecipitation experiments were repeated at least three times. Selected blots were quantified with ImageJ software.

### Real-time PCR

Total RNA was isolated from colonic tissue (that had been snap-frozen in liquid nitrogen, and collected at 24 h) using the RNAsimple Total RNA Kit (Tiangen Biotech, Beijing, China) as described in the manufacturer’s protocol. The RNA concentrations were quantified at 260 nm, and their purity and integrity were determined using a Nano Drop. Reverse transcription and real-time PCR assays were performed to quantify steady-state messenger RNA levels of TLR4, MyD88 and NF-κB at 48 h following CLP (Sham n = 6; CLP n = 6; FMT n = 6). Complementary DNA was synthesized from 0.1μg of total RNA. The following cycling protocol was used: denaturation at 95°C (15 min) and 40 cycles of 95°C (10 sec), 60°C (32 sec). The reporter dye emission (SYBR green) was detected by an automated sequence detector combined with ABI Prism 7500 Real Time PCR System (Applied Biosystems, Foster City, CA).

### Microbiome evaluation

Fecal samples were collected at the time mice were euthanized and frozen until DNA extraction samples were sent to the Shanghai Personal Biotechnology Co., Ltd., where bacterial DNA was extracted. PCR amplification of the bacterial 16S rRNA genes V3-V4 region was performed using the forward primer 338F (5’-ACTCCTACGGGAGGCAGCA-3’) and the reverse primer 806R (5’-GGACTACHVGGGTWTCTAAT-3’). The Illumina MiSeq sequencing platform was amplified (Illumina, San Diego, CA, USA) using the NovaSeq-PE250 sequencing strategy. Sequence denoising or OUT clustering was performed according to QIIME2 dada 2 analysis process or Vsearch software analysis process. The resulting sequences were clustered with 100% similarity against the GreenGenes to database (v. 13.8; https://greengenes.secondgenome.com/)^5^to assign bacterial operational taxonomic units. Alpha and beta diversity comparisons were performed using QIIME2 and taxonomic summaries were generated with QIIME2. Comparison of sample compositions and identification of statistically significant differences were performed with LEfSe using the correction for independent comparisons.

### Statistical analysis

The Kaplan-Meier estimator was used to draw the survival curve of the mice, and the log-rank method was used to compare the survival rates between different groups. The measurement data had a normal distribution, the variance was uniform, and the one-way ANOVA and LSD tests were used for comparison between multiple groups. The Student’s T-test was used to compare the two independent groups, and the measurement result was expressed as the mean ± standard deviation (mean ± sd.). The Kruskal-Wallis *H* test was used for data with non-normal distribution and/or uneven variance. The results were expressed in medians (interquartile range). SPSS 21.0, GraphPad Prism 8.0, Photoshop CS5, Image-Pro Plus, and ImageJ were used for data analysis, and *P* < 0.05 was considered statistically significant.

## Results

### Mortality among three groups

The survival rates of the Sham group, CLP group, and FMT group were compared 7 days following sepsis modeling. The CLP group had a mortality rate of 30% at 24 h and a 50% survival rate at 7 days. There were no deaths at 24 h in the FMT group, and a 90% survival at 7 days, and no deaths in the Sham group. Compared with the Sham group and the FMT group, the mortality of the CLP group was significantly higher, *P* < 0.05. (Figure 1. I).

**Figure 1.**
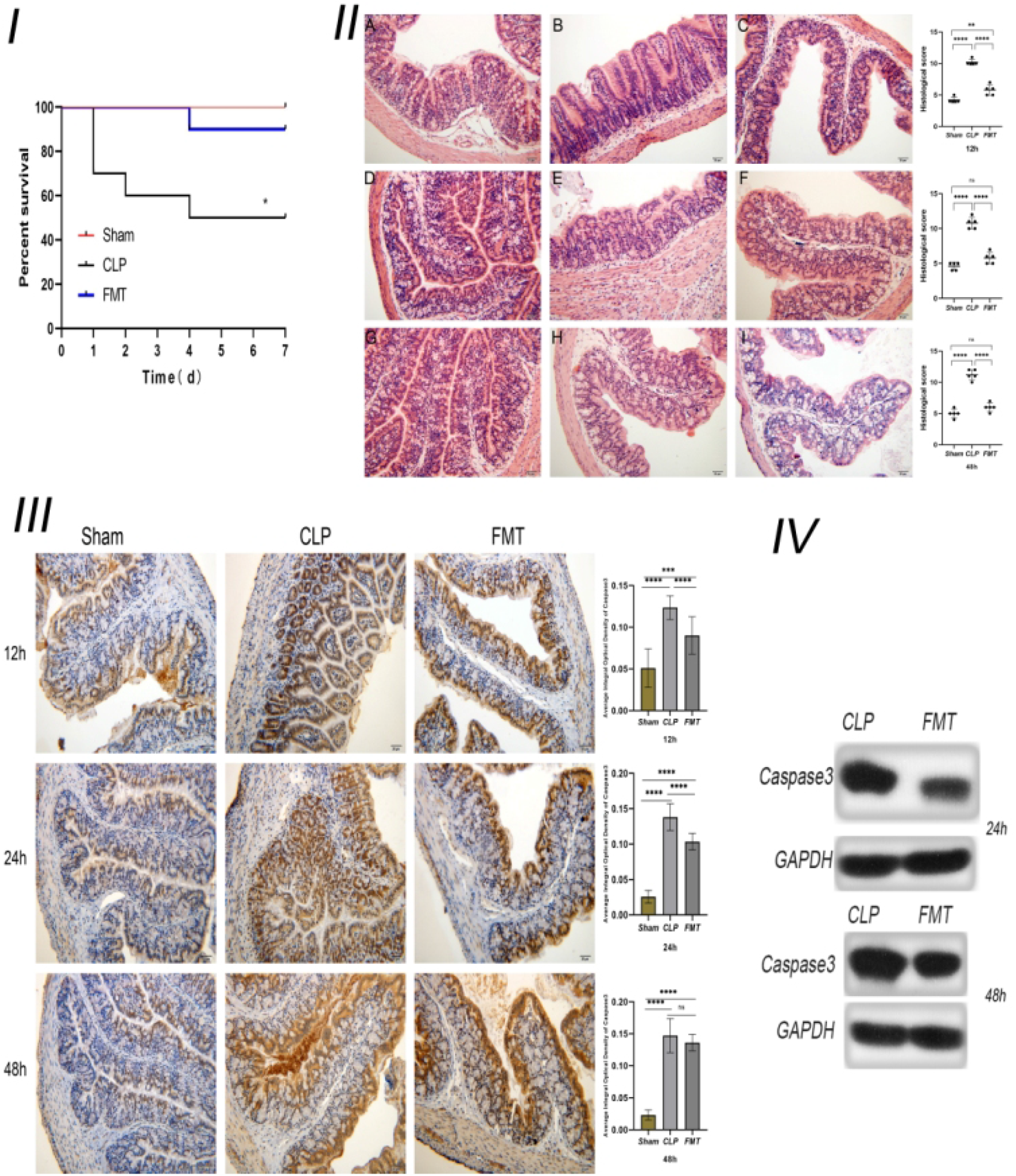
Survival analysis, colonic morphology, and caspase 3 expression among the Sham, CLP, and FMT groups. (I) Seven-day mortality observations in the Sham (red), CLP (black), and FMT groups (blue). There were significant differences between the CLP group and the other two groups, *P* < 0.05, but there was no significant difference between the FMT group and the Sham group; (II) Colonic hematoxylin-ecosin staining of the Sham, CLP, and FMT groups, and their histological scores at 12, 24, and 48 h observed and recorded under a light microscope (20 × 10). A, D, and G show Sham results at 12, 24, and 48 h; B, E, and H show CLP results at 12, 24, and 48 h; C, F, and I show FMT results at 12, 24, and 48 h. The scatter plots on the right side represent the comparison of histological scores between the three groups at 12, 24, and 48 h. (III) Immunohistochemistry of the colon (under a light microscope (20 × 10)) and the average integral optical density of caspase 3 among the Sham, CLP, and FMT groups at 12, 24, and 48 h. (IV) Relative expression of caspase 3 compared with GAPDH between the CLP and FMT groups at 24 and 48 h. * *P* < 0.05, \*\**P* < 0.01, \*\*\**P* < 0.001, \*\*\*\**P* < 0.0001, ns - no statistical difference, respectively.

### Colonic pathology score and apoptosis

Histological scores were performed by professionals who were unfamiliar with our experiment according to the standards proposed by Li et al^23^. The colonic pathology score of the CLP group was significantly higher than the scores of both the Sham and FMT groups at 12, 24, or 48 h, *P* < 0.0001. The colonic pathology score of the FMT group was higher than that of the Sham group at 12 h, *P* < 0.004, but there was no significant difference between the Sham and FMT groups at 24 and 48 h (Figure 1.II). Examination of colonic pathology showed the damage in the FMT group was -significantly less than that in the CLP group, so we next studied the expression of caspase 3 for further verification. The number of apoptotic cells in the CLP group was significantly higher than that of the other two groups at 12 and 24 h, but the average integral optical density (OD) of caspase 3 in the FMT group was close to that of the CLP group at 48 h (Figure 1.III). Therefore, we performed a quantitative analysis of caspase 3 protein. The results of the western blot showed that the expression of caspase 3 in the CLP group at 24 and 48 h was higher than that in the FMT group, *P* < 0.001(Figure 1. IV).

### Mouse serum inflammatory factors TNF-α, IL-6, and IL-10

The concentration in pg/mL of serum IL-6 (91.62 ± 25.53) and IL-10 (7.19 ± 2.40) in the FMT group at 12 h after sepsis modeling were significantly higher than those in the Sham group (IL-6 59.11 ± 12.96; IL-10 3.55 ± 0.72; *P* < 0.05), and there was no significant difference in TNF-α levels (pg/mL) among the three groups.

The serum IL-6 level (130.66 ± 22.52) in the CLP group was significantly higher than that in the Sham group (51.61 ± 5.22) and the FMT group (69.99 ± 16.73; *P* < 0.001) at 24 h. The TNF-α levels in both the CLP (82.97 ± 14.45) and FMT (61.45 ± 16.54) groups were higher than that in the Sham group (27.05 ± 9.44) (*P* < 0.001 and *P* < 0.01, respectively). The IL-10 level in the CLP group (2.74 ± 0.61) was lower than that in the Sham group (7.29 ± 1.12) and the FMT group (8.63 ± 1.24).

The serum IL-6 level in the CLP group (73.82 ± 14.20) was higher than that in the Sham group (43.69 ± 19.02) at 48 h. The TNF-α level in the CLP group (103.53 ± 13.01) was higher than that in the Sham group (24.60 ± 8.89) and FMT group (45.87 ± 9.30) (*P* < 0.001). The TNF-α level in the FMT group was higher than that in the Sham group, *P* < 0.05. There was no significant difference among the three groups in the IL-10 level at 48 h.

All the results are shown in Table 1, Table 2, and Table 3.

**Table 1.**
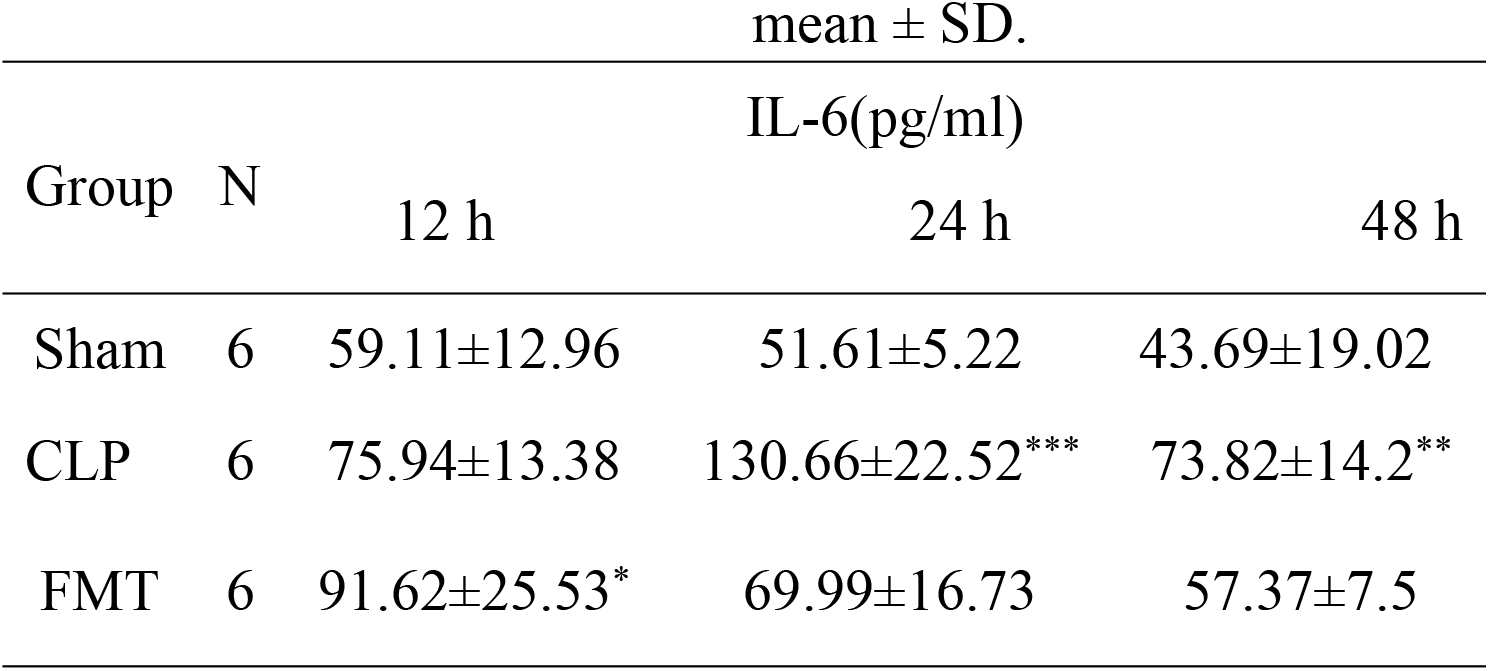
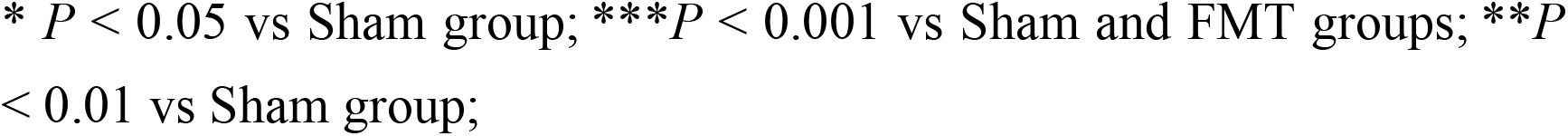
Changes in IL-6 among the three groups. Values are expressed as mean ± SD.

| Group | N | IL-6(pg/ml) |  |  |
| --- | --- | --- | --- | --- |
|  |  | 12 h | 24 h | 48 h |
| Sham | 6 | 59.11±12.96 | 51.61±5.22 | 43.69±19.02 |
| CLP | 6 | 75.94±13.38 | 130.66±22.52*** | 73.82±14.2** |
| FMT | 6 | 91.62±25.53* | 69.99±16.73 | 57.37±7.5 |

\* $P < 0.05$ vs Sham group; \*\*\* $P < 0.001$ vs Sham and FMT groups; \*\* $P < 0.01$ vs Sham group;

**Table 2.**
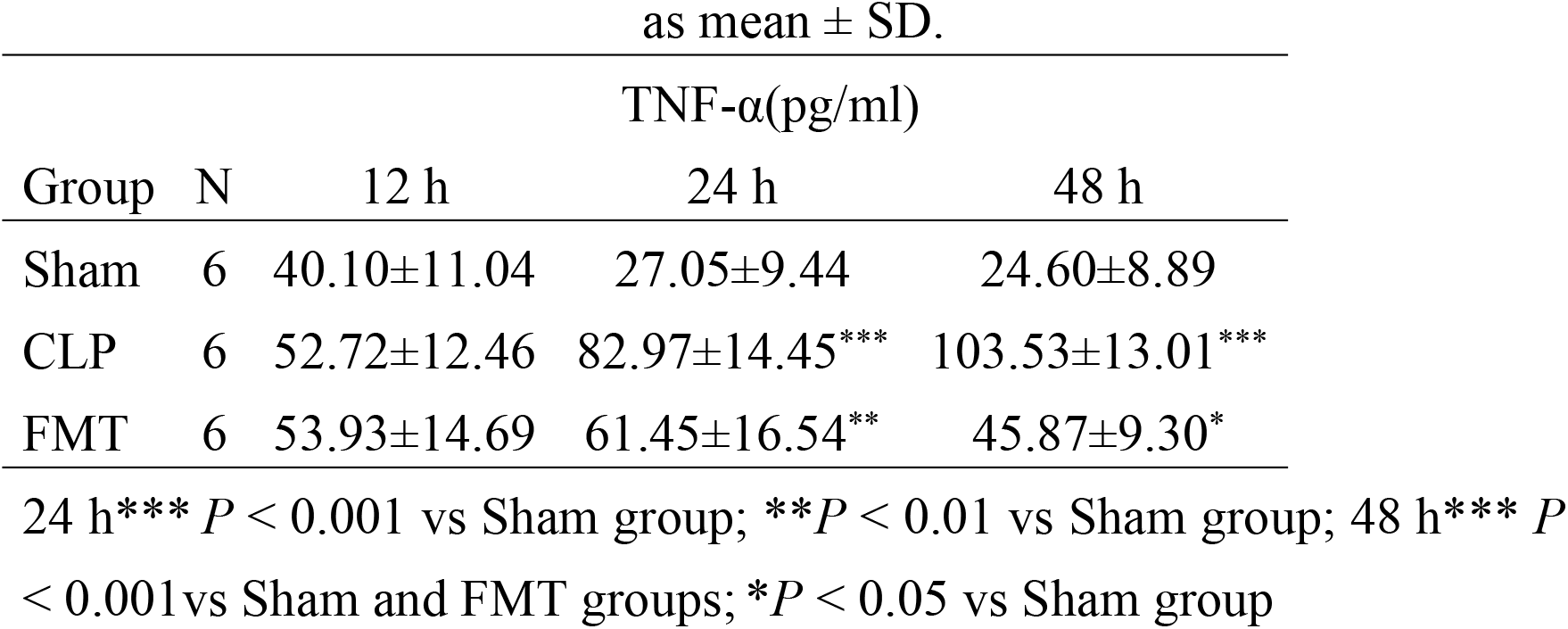
Changes in TNF-α among the three groups. Values are expressed as mean ± SD.

**Table 3.** Changes in IL-10 among the three groups. Values are expressed as mean ± SD.

| Group | N | IL-10(pg/ml) |  |  |
| --- | --- | --- | --- | --- |
|  |  | 12 h | 24 h | 48 h |
| Sham | 6 | 3.55±0.72 | 7.29±1.12 | 5.01±2.15 |
| CLP | 6 | 5.04±1.51 | 2.74±0.61*** | 4.3±1.74 |
| FMT | 6 | 7.19±2.40* | 8.63±1.24 | 6.20±1.07 |

\* $P < 0.05$ vs Sham group; \*\*\* $P < 0.001$ vs Sham and FMT groups.

The expression of IL-6 in the CLP group was the highest at 24 h after modeling, and then decreased, but it was still higher than that in the Sham group at 48 h. Expression of IL-6 in the FMT group was slightly higher than that in the CLP group at 12 h, but there was no significant difference, to the contrary, IL-6 levels were markedly lower than the CLP group at 24 and 48 h. The TNF-α level in the CLP group continued to increase, and it was the highest among the three groups at 48 h, however, the IL-10 level was lowest at 24 h after modeling (Figure 2).

**Figure 2.**
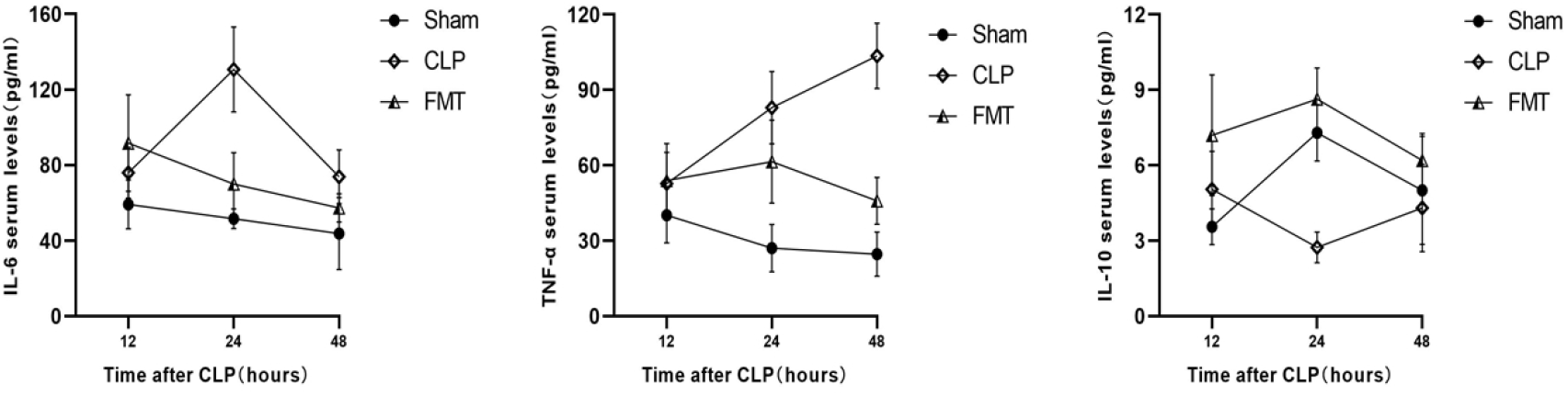
The serum IL-6, TNF-α, and IL-10 among the Sham, CLP and FMT groups at 12, 24, and 48 h.

### The thickness of the mucus layer (nm) and MUC2 expression

The AB-PAS method was used to detect the colonic mucus layer thickness at 12, 24, and 48 h after sepsis modeling in the three groups. The mucus layer thickness of mice in the CLP group (2248.864 ± 603.939, 2046.108± 664.865, and 1806.371 ± 579.875, at 12, 24, and 48 h respectively) was significantly lower than that of the Sham group during the same timepoint, *P* < 0.0001. The thickness of the mucus layer in the FMT group was significantly greater than that in the CLP group, *P* < 0.01. Compared with the Sham group, the thickness of the mucus layer in the FMT group had no difference at 24 h, *P* = 0.4473, but had a significant difference at 12 and 48 h, *P* < 0.0001, and *P* < 0.05, respectively (Figure 3. I). The fluorescence expressions of MUC2 at 12, 24, and 48 h in the three groups were observed by microscope and recorded, and the gray value of the green channel was calculated and statistically analyzed. Excepting that the expression levels of MUC2 in the Sham group and the FMT group were not statistically different at 12 h, there was a significant difference between them at 24 and 48 h, *P* < 0.0001. The expression of MUC2 in the CLP group at 12, 24, and 48 h was lower than that of the other two groups (Table 1, Figure 3. II).

**Figure 3.**
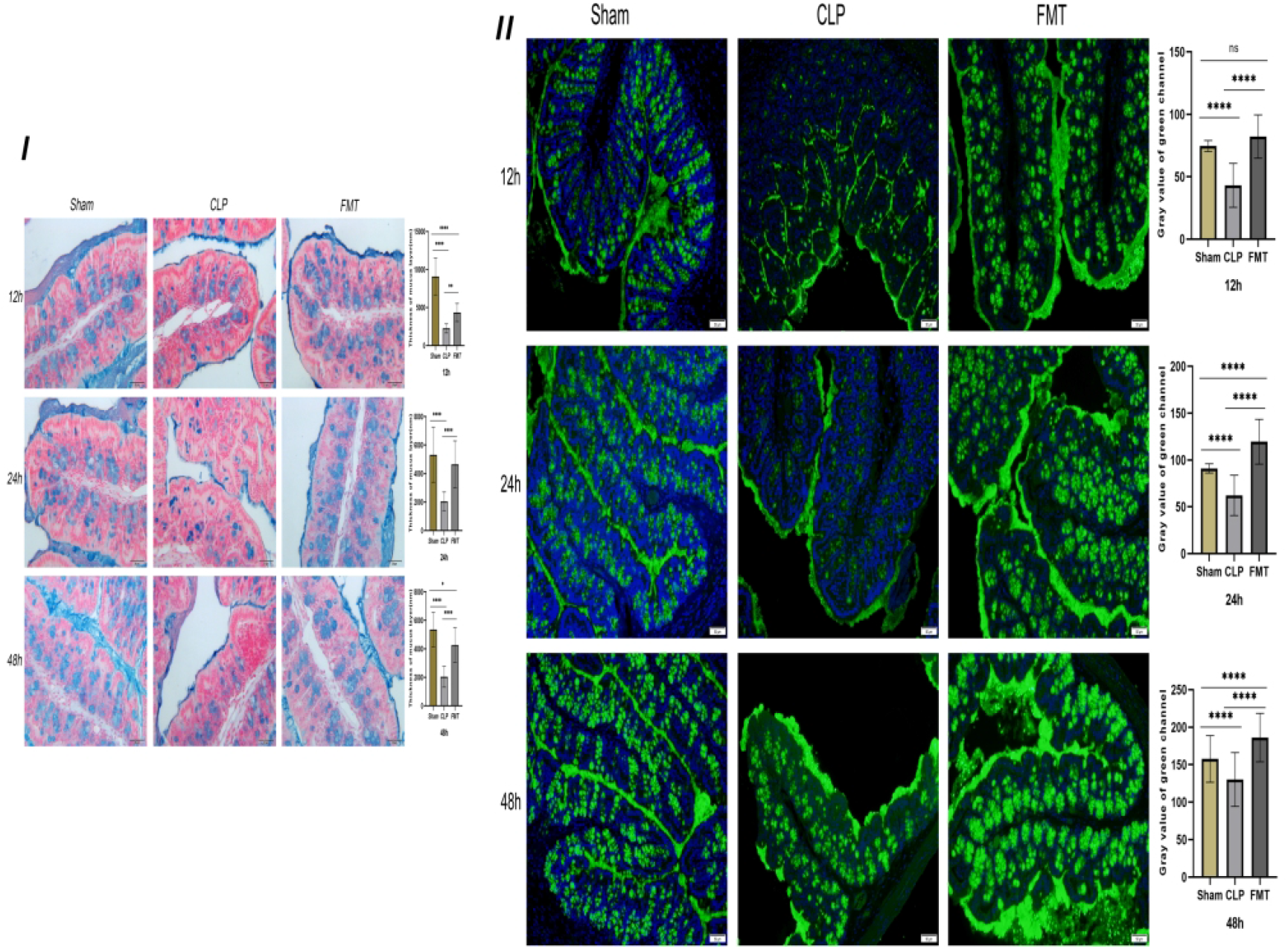
The mucus layer thickness and MUC2 expression in the Sham, CLP, and FMT groups at 12, 24, and 48 h. (I)The AB-PAS method was used to measure the thickness of the mucus layer in the Sham, CLP, and FMT groups at 12, 24, and 48 h under a light microscope (20 × 10). The blue color indicates goblet cell secretion and mucus layer. The histogram on the right shows the statistical results of the comparison of the three groups. There was no significant difference between the Sham group and the FMT group in mucus layer thickness at 24 h. (II) MUC2 expression (green fluorescence) was observed in the three groups at 12, 24, and 48 h. The histogram shows the statistical results of all three groups. There was no significant difference between the Sham group and the FMT group at 12 h. \**P* < 0.05, \*\**P* < 0.01, \*\*\*\**P* <0.0001, ns : no significant difference.

**Figure 4.**
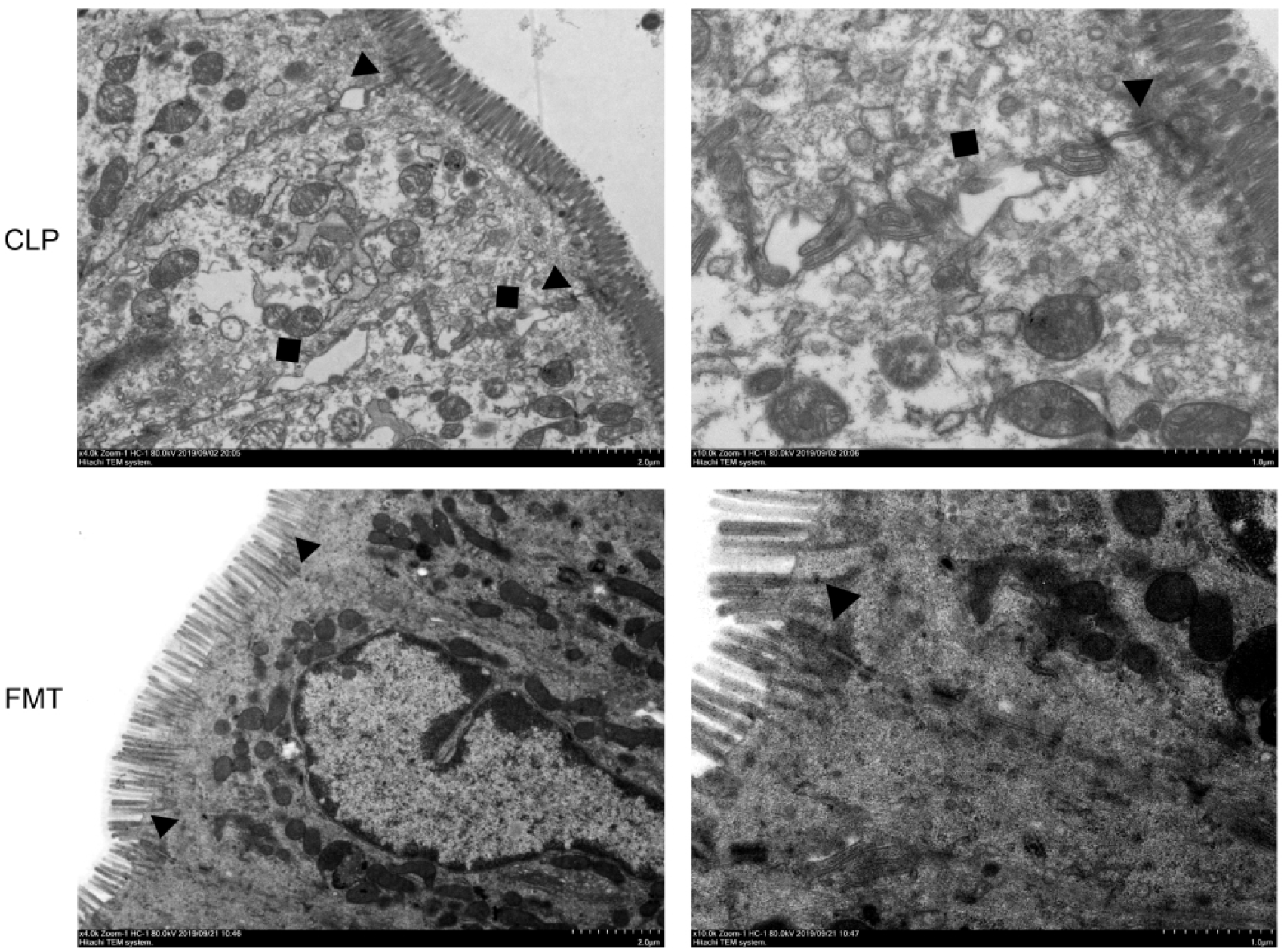
Intestinal epithelial cell junctions. Intestinal epithelial cell junctions in the CLP group and FMT group at 48 h. The left image was enlarged 4K, the right one was enlarged 10K. ▲Tight junction and intermediate junction between intestinal epithelial cells ■The space between intestinal epithelial cells.

### Transmission electron microscope

Intestinal epithelial cells and intracytoplasmic organelles in the CLP group were significantly swollen, and the microvilli were arranged neatly, with uniform thickness and partial shedding. The tight junctions, and the structure of the intermediate junctions were blurred, and the gap between junctions was slightly widened in the CLP group. Portions of the desmosomes and tension wires had disappeared. The intercellular space was widened and the mitochondria had swelled. Intestinal epithelial cells and intracytoplasmic organelles in the FMT group were slightly swollen. The microvilli were arranged neatly and uniformly in thickness, and the local area was slightly detached. The tight junctions between epithelial cells, and the structure of the intermediate junctions were fuzzy, and the gap was slightly widened in the FMT group. The number of desmosomes was slightly reduced, the surrounding tension filaments were abundant, and the mitochondria were slightly swollen.

### Tight junction proteins

We compared the average integral OD of occludin between the CLP and the FMT groups at 12, 24, and 48 h. The results showed that occludin expression in the CLP group at 12, 24, and 48 h was significantly lower than expression in the FMT group (Figure 5. I). We further verified occludin and ZO-1 protein expression by western blot test. The results showed the relative expression of these two proteins in the CLP group was significantly lower than that in the other two groups at 24 or 48 h, (*P* < 0.001). The relative expression of the two proteins was highest in the Sham group, intemediate in the FMT group and lowest in the CLP group (Figure 5. II).

**Figure 5.**
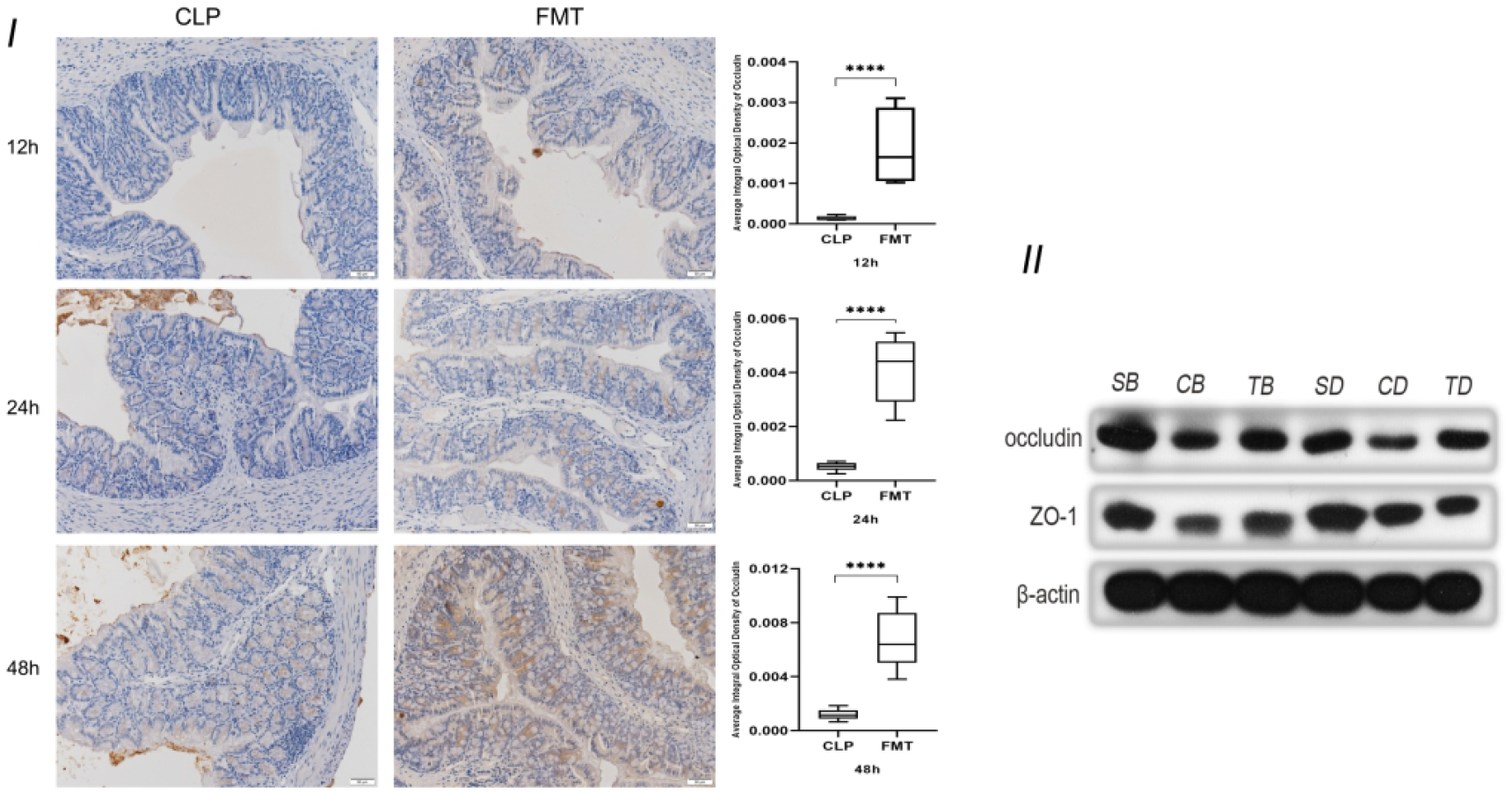
Comparison of tight junction protein expression (I)Comparison of occluding expression in the CLP and FMT groups at 12, 24 add 48 h, respectively, and observed under a light microscope (20 × 10). Significant results are shown in the box plot on the right, \*\*\*\**P* < 0.0001; (II) Relative expression of occludin and ZO-1 compared with β-actin expression in the Sham, CLP and FMT groups at 24 or 48 h. SB, CB, TB, SD, CD, and TD represent the 24 h Sham, CLP, and FMT groups and 48 h Sham, CLP, and FMT groups, respectively.

### TLR4, MyD88, and NF-κB protein levels and mRNA levels

We analyzed and compared the expression of TLR4, MyD88, and NF-κB relative to GAPDH at 24 and 48 h in the three groups. Except that there was no difference in the expression of TLR4 protein between the Sham and FMT groups, expression levels of the other two proteins in the three groups were significantly different compared with each other. Expression of the three proteins in the CLP group at 24 and 48 h was significantly higher than those in both the Sham and the FMT groups at the same timepoints (*P* < 0.05)(Figure 6. I). Coincidently, mRNA expression trends in the three groups were similar to the trends in protein expression. The relative expressions of TLR4 (2.161 (1.971)), MyD88 (1.577 (2.552)), and NF-κB (1.489 (1.447)), in the CLP group were significantly higher than those in the Sham and FMT groups. Expression of MyD88, and NF-κB had a significant difference in the CLP group compared with the Sham and FMT groups, *P* < 0.05. There was no difference in the expression of the three indicators between the Sham group and the FMT group (Figure 6. II).

**Figure 6.**
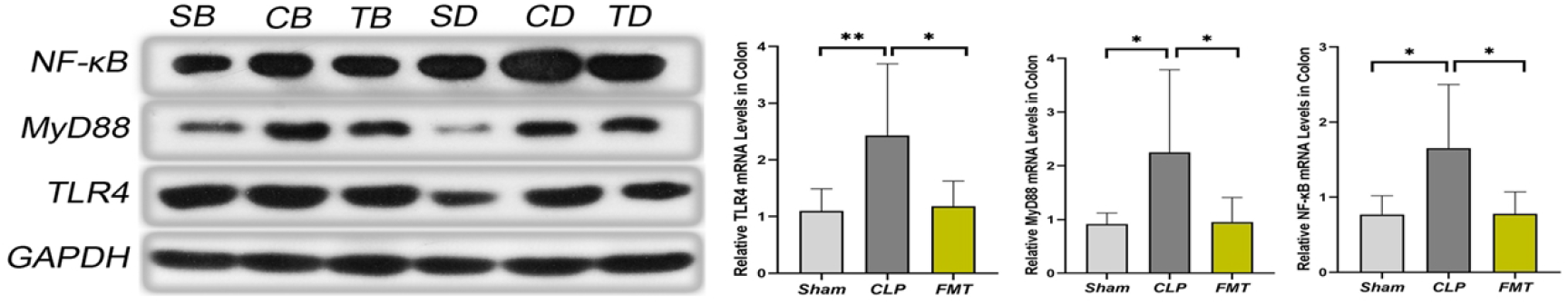
TLR4, MyD88, and NF-κB expression and mRNA levels (I) Expression of TLR4, MyD88, and NF-κB compared to GAPDH protein among Sham, CLP, and FMT groups at 24 and 48 h. SB, CB, TB, SD, CD, and TD represent 24 h Sham, CLP, and FMT groups and 48 h Sham, CLP, and FMT groups, respectively. (II) Relative TLR4, MyD88, and NF-κB levels in the colon. \**P* < 0.05, \*\**P* < 0.01.

### 16SrRNA sequence analysis

Firmicutes and Bacteroidetes were the main bacteria, accounting for 55% and 33% of the abundance found in the normal mice, respectively. The Sham group was dominated by Firmicutes, at 66%. Proteobacteria was the dominant bacteria in the CLP group. The relative abundance of Proteobacteria observed in the mortality observation of the CLP groups following sepsis modeling at 12 h, 24 h, and 7-day were 48%, 66%, and 42%, respectively. The relative abundance of unclassified bacteria in the CLP group at 48 h and the 7-day mortality observation in the FMT group was 70% and 41%. The FMT group at 24 and 48 h were dominated by Firmicutes and Bacteroidetes, accounting for 48%, 27%, and 37%, 26%, and the composition closely resembled the composition found in mice. The main bacteria in the FMT group at 12 h were Proteobacteria and Verrucomicrobiae, with a relative abundance of 33% and 39%, respectively. Verrucomicrobiae accounted for 22% in the CLP group at 24 h. (Figure 7. I). The 12 h Sham group had a similar fecal richness and diversity compared with normal mice. The fecal richness and diversity were significantly lower in the CLP group at 12, 24, and 48 h and 7-day mortality than those of normal mice. We observed that the fecal richness and diversity were lowest at 48 h in the CLP group, but there was a slight recovery in the 7-day still alive mice in the CLP group, a significant difference compared with the normal mice. The richness and diversity of the flora in the FMT group were higher than what was observed in the CLP group at all time points, and there was more fecal diversity than the normal mice at 24 h in the FMT group. There was no significant difference in fecal richness and diversity in the 7-day mortality FMT mice compared with the CLP group at the same timepoint (Figure 7. II). The dimensionality reduction of multi-dimensional microbial data was performed through NMDS analysis, and the main trends of data changes were displayed through the distribution of samples on a continuous sorting axis. The data were also classified by cluster analysis. In the NMDS analysis, the clusters within the group were well and the difference between the groups was large (Figure 7. III). LEfSe analysis can directly perform simultaneous difference analysis on all classification levels, and at the same time, it emphasizes finding robust differences between groups, that is, marker species. The main species found in normal mice were Firmicutes and Bacteroidete. The species with high levels in the Sham group were Firmicutes and Lactobacillus, while the differential species in the CLP group were Bacillales and Staphylococcaceae at 12 h, Enterobacteriales and Proteobacteria at 24 h, Planococcaceae at 48 h, and Deltaproteobacteria, Desulfovibrionales, and Erysipelatoclostridium at 7-day mortality. The FMT group was modeled with high levels of bacteria, including Verrucomicrobiae, Akkermansia, and Ruminococcus at 12 h, Lachnospiraceae group, Bifidobacteriales, Actinobacteria at 24 h, and Burkholderiaceae, Bacteroides, and Butyricimonas at 48 h (Figure 7. IV). The core of the KEGG database is a biological metabolic pathway analysis database, in which metabolic pathways are classified into six categories, including metabolism, genetic information processing, environmental information processing, cellular processes, organismal systems, and human diseases. We found a higher relative abundance in infectious diseases. (Figure 7. V). Therefore, we further analyzed the species composition of the differential pathways, and found that the Lachnospiraceae contributed the most to L-lysine fermentation to acetate and butanoate (Figure 7. VI).

**Figure 7.**
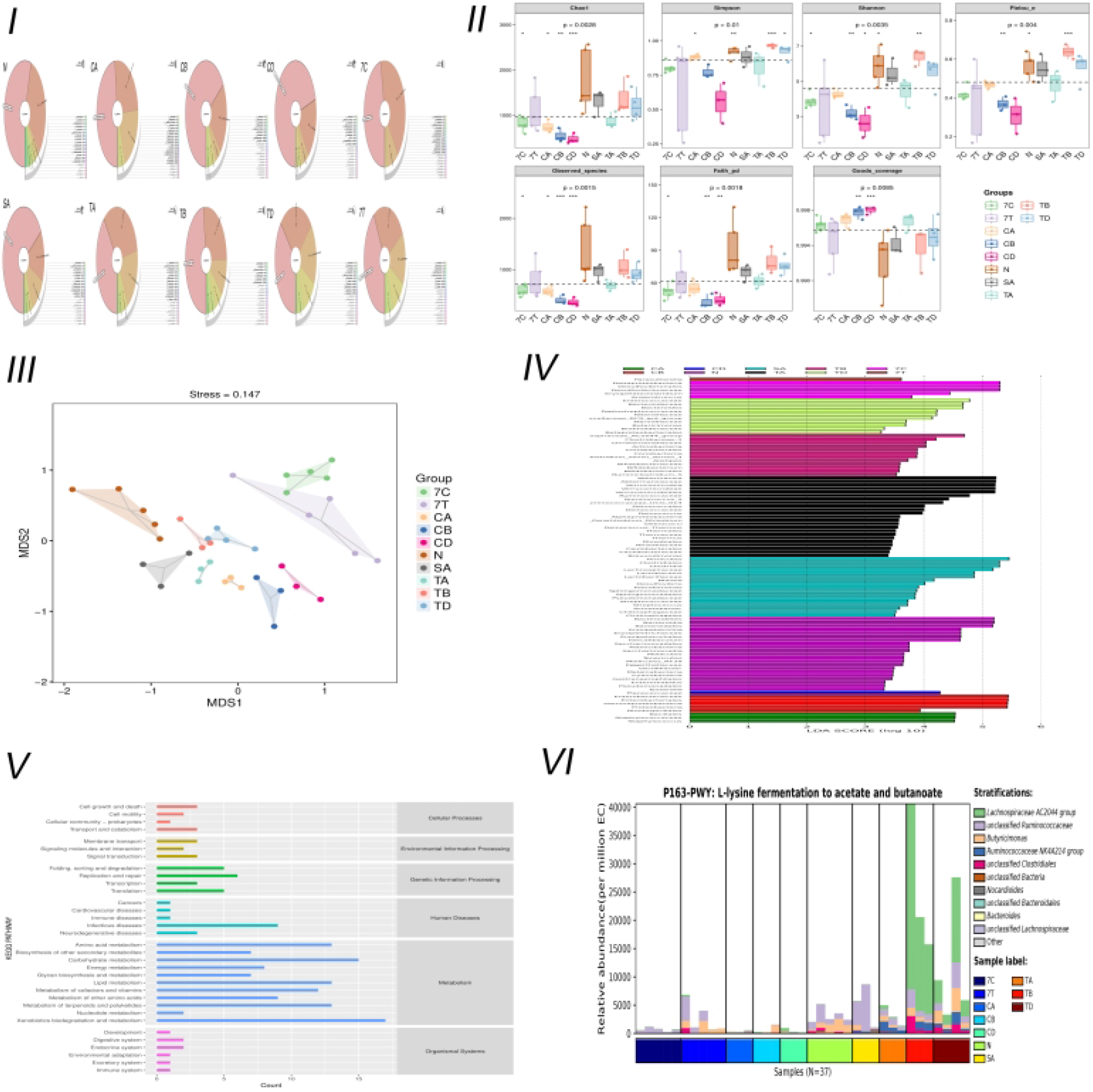
Microbiota analysis. (I) Krona species at phylum among the Normal, Sham, CLP, and FMT groups. CA, CB, CD, and TA, TB, TD represent the 12, 24, 48 h Sham, and 12, 24, and 48 h FMT groups, respectively. N-normal mice. SA represents the 12 h Sham group. (II) Alpha diversity. (III) NMDS two-dimensional sorting diagram. The closer the distance between the two points in the figure, the smaller the difference between the microbial communities in the two samples. (IV) LEfSe (LAD Effect Size). A longer bar denotes a more significant difference in the taxon. The color of the bar graph indicates the most abundant sample group corresponding to the taxon. (V) Predicted abundance map of KEGG secondary functional pathways. (VI) Species composition map of differential MetaCys metabolic pathways. The abscissa shows different groups. The order of the samples in the groups was sorted according to the similarity of the data; the ordinate was the relative abundance of the metabolic pathways, and the species different levels of contribution to the metabolic pathways were displayed in different colors at different levels. (N = normal mice, 7C = CLP group mortality observation, 7T = FMT group mortality observation, SA = Sham group 12 h; CA, CB, and CD = CLP group at 12, 24, and 48 h timepoints, respectively, and TA, TB, and TD = FMT group at 12, 24, and 48 h timepoints).

### Analysis of short-chain fatty acids (SCFAs) by LC-MS

We detected the content of short-chain fatty acids, including acetate, propionate, isobutyrate, butyrate, valerate, isovalerate, caproate, and heptanoic acid in the CLP group (CD) and the FMT group (TD) at 48 h. The first four acids were the most important SCFAs, therefore, we used these four major short-chain fatty acids for PCA analysis. (n = 7 per group) (Figure 8.).

**Figure 8.**
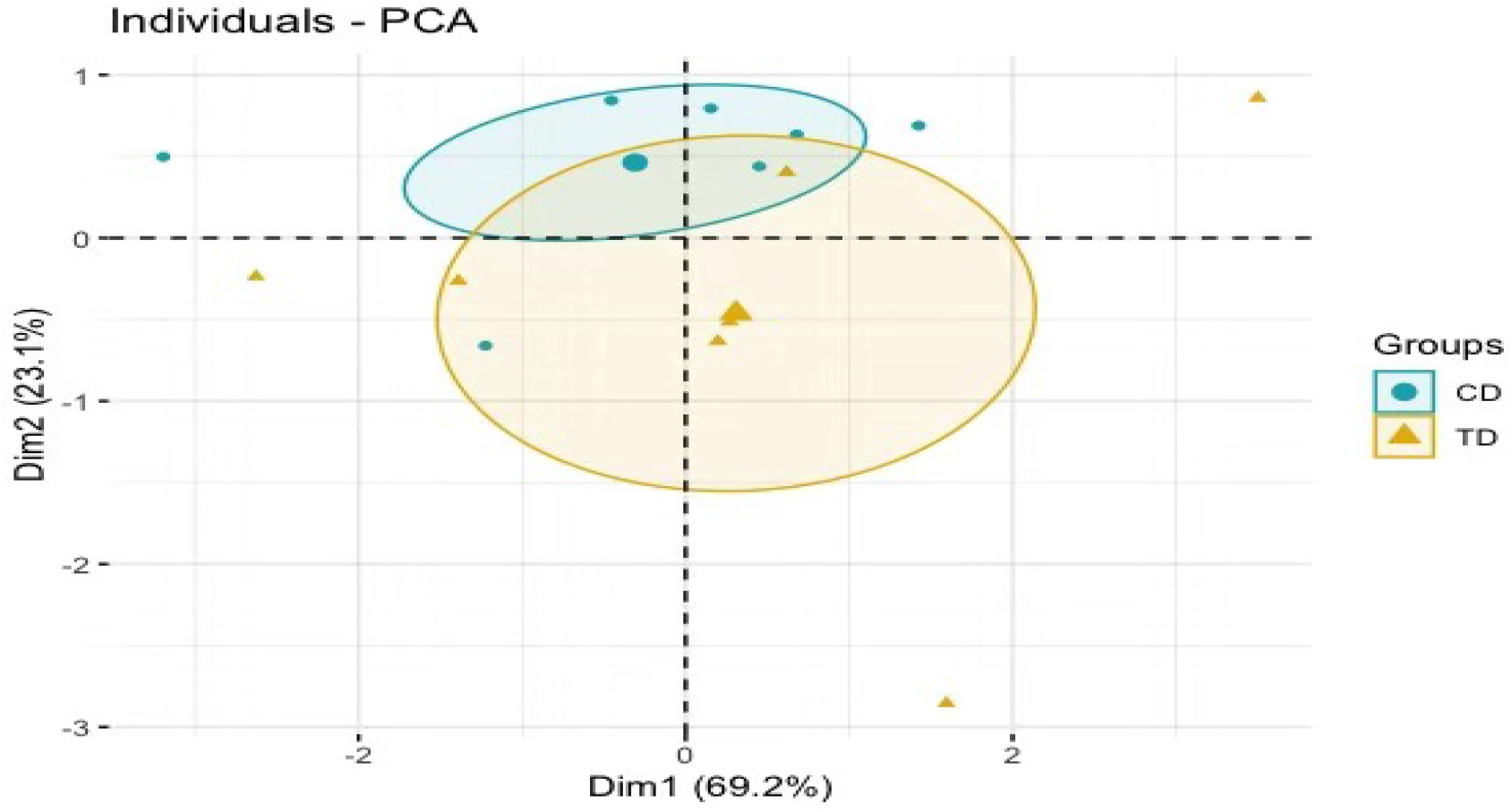
Analysis of four major short-chain fatty acids in the CD and TD groups.

## Discussion

The human gastrointestinal (GI) tract is home to over 10^14^ microorganisms comprising bacteria, archaea, viruses, yeast, fungi, and phages, collectively designated as GM^27–29^. The four phyla dominating the adult human microbiome are Firmicutes, Bacteroidetes, Proteobacteria, and Actinobacteria. Of those, 90% of all of the intestinal bacterial species fall under the most heavily studied: Firmicutes and Bacteroidetes^13^. Firmicutes are Gram-positive bacterial taxa composed of many familiar-sounding genera including Clostridia, Streptococcaceae, Staphylococcaceae, Enterococcaceae, and Lactobacillae, many of which are facultative anaerobes. Bacteroidetes, on the other hand, are Gram-negative bacteria composed largely of Bacteroides species, which are obligate anaerobes^30^. Fecal microbiota transplantation refers to the transplantation of functional bacteria from the feces of healthy donors into the patient’s GI tract to restore the intestinal microecological balance and subsequently treat diseases related to microbial imbalances^17, 31^. This study was an attempt to explore whether FMT can maintain the integrity of the intestinal flora and protect the intestinal barrier function in mice with sepsis. We found that there was a flora imbalance 12 h after the modeling of sepsis. Proteobacteria had an absolute advantage, but Firmicutes and Bacteroidetes were reduced, which was consistent with previous studies^12^. The relative abundance of Firmicutes and Bacteroidetes recovered slightly over time, but Proteobacteria in the 7-day mortality CLP group still accounted for 42% of intestinal flora, which was significantly higher than that of normal mice. Analysis of the bacterial flora in the FMT group revealed that Firmicutes and Bacteroidetes were most prevalent and the relative abundance of Proteobacteria was low, a bacterial composition similar to that of normal mice. The Alphaproteobacterial load was increased and the Betaproteobacterial load was diverse in the FMT group. All of these observations indicate that the intestinal flora of septic mice had been replenished after FMT. This is similar to the previous analysis of FMT use in recurrent *C. difficile* infection and inflammatory bowel disease. It is believed that FMT can maintain the intestinal bacterial balance by increasing the diversity of the flora and restoring intestinal flora after external interference^32^. However, in the 7-day mortality observation FMT group, the richness and diversity of bacterial flora were lower compared with the FMT group at 24 and 48 h. As aforementioned, the dose and the number of infusions of FMT may have contributed to this result^32^. This experiment confirmed that the 7-day mortality rate of septic mice was significantly higher than that of the FMT group. Early application of FMT can effectively reduce the mortality of septic mice, most likely due to the reconstruction of intestinal flora.

Gut barrier dysfunction can be considered both a result and a cause of sepsis development^19^. There are four mechanisms associated with gut barrier failure. The first, most common phenomenon, is a loss of enterocyte integrity; the second, an increased paracellular permeability; the third, a breakdown of the mucus layer; and the fourth, a decreased mucosal immunity^33^. Therefore, the interaction between the microbiome and the intestinal immune system is critical for maintaining mucosal homeostasis^15^.

Tight junctions play an important role in maintaining the integrity of the mucosal epithelium^34, 35^, and it is generally defined as life-threatening organ failure in the setting of critical illness^36, 37^. The barrier function is mediated by changes in the tight junctions and associated proteins and allows outflow of luminal contents and likely damages distant organs, which may contribute to MODS^12^. It has been shown that TNF-α, IL-1β, and IL-6 cause an increase in intestinal tight junction (TJ) permeability and contribute to the inflammatory process by allowing luminal antigenic penetration. Conversely, the anti-inflammatory cytokine IL-10 was shown to promote intestinal TJ barrier function^38, 39^. Critical illness induces hyper-permeability of the gut barrier which begins as early as 1 hour after the onset of sepsis and lasts at least 48 h^12, 40^. We compared transmission electron microscopy results at 48 h between the CLP and FMT groups. The TJs of the septic mice were blurred more than those of the FMT mice. The cell gap was significantly wider, and both cells and organelles were swollen. Mice had not only intestinal barrier dysfunction but also cellular dysfunction. The main functions of occludin are regulation and sealing of TJs^41, 42^. That ZO-1 outperformed other TJ markers may reflect the concomitant organ epithelial injury which occurs in MODS^36^. Therefore, we observed the expression of the above two proteins at 24 and 48 h in the three groups, and the results showed that occludin and ZO-1 in the CLP group were significantly lower than those in the Sham group and the FMT group. Levels of the two indicators in the FMT group were higher than those in the CLP group, or they were similar to those in the Sham group.

The intestinal mucus layer is divided into two layers, a dense inner layer and a loose outer layer^26, 43^. The mucous gel is formed by high molecular-weight mucins secreted by the goblet cells and the outer mucus layer serves as a habitat for the resident microflora by providing a beneficial microenvironment^43, 44^. The amount and composition of the mucus layer reflect the balance between mucus secretion, and its erosion and degradation by bacteria^45^. Research has shown that the thickness of the mucus layer is dependent on commensal bacteria^12^. We performed blinded thickness measurements of the colonic mucus layer in each mouse using Alcian blue-stained sections. We further validated the thickness of the mucus layer by immunofluorescence staining of the MUC2 mucins using a-MUC2 antibody^25^, and observed that the thickness of the mucus layer in septic mice was significantly reduced, and the thickness in the FMT group was significantly increased. This change was consistent with the changes in the flora of the two groups.

Although commensal bacteria reside innocuously in the gut mucosa, they share common microbe-associated molecular patterns (MAMPs) with the pathogenic bacteria that invade through the intestinal epithelium^46, 47^. Commensal bacteria, therefore, have the potential to activate immune responses through pattern recognition receptors such as toll-like receptors (TLRs) and nucleotide-oligomerization domain (NOD)-like receptors (NLRs)^47^. Mild or physiologically acceptable levels of stimulatory signals provided by commensal flora are essential for the development and maintenance of an appropriate mucosal immune system, which provides the first line of defense together with the epithelial barrier system against undesired, foreign antigens^44^.

TLRs are type I transmembrane glycoproteins and serve as signal transduction receptors for innate immunity and inflammation^15^. TLR4 is the best-characterized pathogen-recognition receptor. Its downstream effects are varied, and the TLR/MyD88/p38 MAPK/NF-κB pathway is popularly believed to play a critical role in the inflammatory response^33, 48^. A major pathophysiological mechanism of sepsis involves recruitment of inflammatory cells and generation of an overwhelming pro-inflammatory response. Cell-wall components from Gram-negative and Gram-positive bacteria activate PRRs such as TLR4 and TLR2, respectively, resulting in a “cytokine storm” of pro-inflammatory mediators generated mainly via the mitogen-activated protein kinase and NF-κB pathways^19^. In this study, the TLR4/MyD88/NF-κB pathway in septic mice increased significantly at both the protein level and the gene level, and the inflammatory factors TNF-α and IL-6 increased at different levels at 24 and 48 h after modeling, while IL-10 decreased at 24 and 48 h after modeling. In contrast, the FMT group showed an increase in inflammatory response inhibition. This is consistent with the pathological score of bowel injury in the three groups. It was observed that the amount of apoptosis protein caspase 3 was different between the CLP group and the FMT group, and the amount of apoptosis in the CLP group was significantly higher than that of the FMT group.

The GM support the development of the metabolic system and the maturation of the intestinal immune system by providing beneficial nutrients, by synthesizing vitamins and short-chain fatty acids (SCFAs)^15^. Acetate, especially, is often considered to play a critical role in protection against pathogenic infection^16, 49^. Studies have shown that SCFAs have anti-inflammatory and immunoregulatory activities and may reduce butyrate-producing bacteria such as Ruminococcaceae, Faecalibacterium, and Roseburia^13^. Therefore, we compared the relative abundance of Ruminococcaceae in each group, and the results suggest that fecal bacteria were significantly reduced in the sepsis model group, while in the FMT group, bacterial counts were similar to or even slightly higher than those found in normal mice. (In our study, the relative abundance of normal mice was 0.07%, 0.0025% in the CLP group and 0.067% in the FMT group at 48 h). It appears that Ruminococcaceae plays a role in the inflammatory response, and fecal microbiota transplantation can effectively improve the intestinal bacteria composition, thereby improving the inflammatory state. We found through functional prediction that the relative abundance of intestinal flora was significantly different in infectious diseases. Further analysis of the species composition of the different pathways revealed that Lachnospiraceae contributed the most to L-lysine fermentation to acetate and butanoate, consistent with previous studies^50^. This indicated that Lachnospiraceae may be the key bacteria for the effectiveness of FMT. We further explored the changes of SCFAs within 48 h in the CLP group and the FMT group. We mainly studied acetate, propionate, isobutyrate and butyrate. After drawing the PCA chart, we found that there may be a different trend between the two groups, but further studies are needed.

This study has certain limitations. First, we selected the C57BL/6 mouse as our animal model, so it may be difficult to generalize our conclusions to humans. Second, we administrated FMT once a day, and the 7-day mortality observation group received FMT once a day for the first three days. We did not attempt to discern the complete reconstruction time of intestinal flora after initiating FMT, nor did we compare the frequency and number of fecal bacteria transplantation, but it appears that early application of FMT has a protective effect on the intestinal function of septic mice. Thirdly, this experimental study does not involve acquired immunity, a topic demanding further experimentation and verification. Fourth, the transmission electron microscope observed that the effect of FMT is not limited to the intercellular connection, but also involves mitigation of organelle damage. We did not explore the reason for this difference. Finally, the issue of metabolics needs deeper study.

## Conclusion

GM imbalance exists early in sepsis. Fecal microbiota transplantation can improve morbidity and effectively reduce mortality in septic mice. In our study, after FMT, the abundance and diversity of the gut flora were restored, and the microbial characteristics of the donors changed. Fecal microbiota transplantation can effectively reduce epithelial cell apoptosis, improve the composition of the mucus layer, upregulate the expression of TJ proteins, and reduce intestinal permeability and the inflammatory response, thus protecting the intestinal barrier function. After FMT, Lachnospiraceae contributes the most to intestinal protection through enhancement of the L-lysine fermentation pathway, resulting in the production of acetate and butanoate, and may be the key bacteria in short-chain fatty acid metabolism that promotes the success of fecal microbiota transplantation.

**Table 4.**
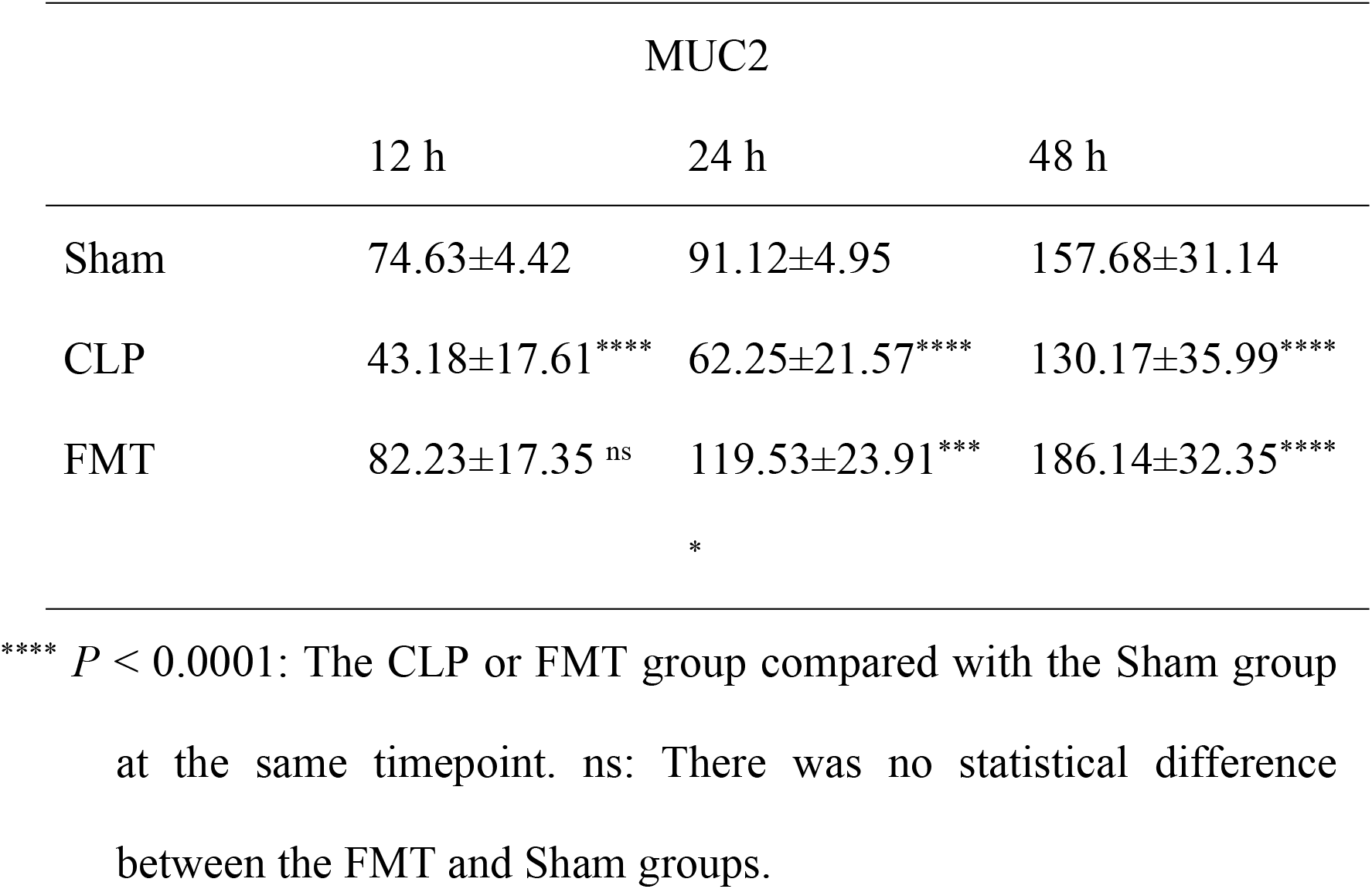
Gray value of the green channel (MUC2 expression), expressed as mean ± SD.

